# The force required to remove tubulin from the microtubule lattice

**DOI:** 10.1101/2022.03.28.486117

**Authors:** Yin-Wei Kuo, Mohammed Mahamdeh, Yazgan Tuna, Jonathon Howard

## Abstract

Severing enzymes and molecular motors extract tubulin from the walls of microtubules by exerting mechanical force on subunits buried in the lattice. However, how much force is needed to remove tubulin from microtubules is unknown, as is the pathway by which subunits are removed. Using a site-specific functionalization method, we applied forces to the C-terminus of α-tubulin with an optical tweezer and found that a force of ∼30 pN is required to extract tubulin from the microtubule wall. Consistent with this force, we show that several kinesins can also extract tubulin. Additionally, we discovered that partial unfolding is an intermediate step in tubulin removal. The unfolding and extraction forces are similar to those generated by AAA-unfoldases, suggesting that severing proteins such as spastin and katanin use an unfoldase mechanism. Our results reveal the response of tubulin to mechanical force and advance our understanding of severing enzymes and microtubule stability.

## Introduction

Microtubules serve as tracks for motor proteins and provide mechanical support to eukaryotic cells ^1^. Eukaryotes encode many microtubule-associated proteins that modulate the dynamics and shapes of the microtubule cytoskeleton to allow cells to move, divide and change shape ^2–5^. Microtubule-severing enzymes (or severases) are AAA-ATPases that can sever microtubule filaments into shorter pieces using the energy of ATP-hydrolysis ^6^. Recent work has demonstrated that the combination of severases, dynamic instability, and microtubule end regulators constitute a versatile mechanism to control the number, length and spatial organization of microtubules ^7–11^, accounting for a wide spectrum of cellular functions that require severing enzymes (reviewed in ^12–14^).

The current model for microtubule severing proposes that severases pull tubulin subunits out of the microtubule lattice by exerting mechanical forces on the C-terminal tails (CTTs) of the tubulin ^15^. The microtubule filament is thought to break when enough tubulin subunits have been removed from the lattice. This model is largely based on sequence and structural similarity to other AAA+ unfoldases and disaggregases such as Hsp100 ^16,17^, ClpX ^18,19^ and Vps4 ^20,21^ that unfold or disassemble proteins (or protein complexes). By using ATP-hydrolysis, these enzymes produce mechanical forces that pull proteins through a central channel made by the six AAA domains ^22^. Biochemical and structural studies show that severing enzymes bind tubulin CTTs ^23–25^, but experimental observations of the force generation and tubulin extraction steps are absent.

In addition to severing enzymes, molecular motors such as kinesins and dynein can also remove tubulin subunits from the shafts of microtubules as they walk ^26–28^. This removal of tubulin dimers from the internal lattice is distinct from the well-known activities of motors removing tubulin from microtubule ends ^29^. Bending of microtubules by fluid flow can also cause dissociation of tubulin subunits from the lattice ^30^. Thus, several lines of evidence suggest that mechanical forces can facilitate the removal of tubulin subunits from the microtubule shaft ^31^. However, direct experimental investigation of the tubulin extraction process is lacking, due, in part, to the difficulty of applying force to a single tubulin subunit in a site-specific manner.

Here, we address two central questions concerning the tubulin extraction process. First, how much force is needed to pull out a tubulin subunit? And second, is tubulin unfolded during the pulling process?

## Results

### Development of site-specific labeling method

To apply forces to tubulin, we developed a strategy to introduce a functional handle specific to the C-terminus of α-tubulin by exploiting the broad substrate tolerance of tubulin-tyrosine ligase (TTL) reported previously ^32,33^. The tubulin CTTs are exposed on the outer surface of microtubule and are the proposed pulling sites of severing enzymes. Thus, tubulin CTTs are suitable handles through which to apply mechanical forces. TTL catalyzes the covalent addition of tyrosine and tyrosine analogs to the C-terminus of detyrosinated α-tubulin. We expressed and purified human TTL with an N-terminal His_6_-SUMO tag (Fig. S1). The tyrosination activity of purified TTL was verified by the increase of tyrosinated tubulin after incubating TTL with bovine brain tubulin in the presence of ATP and L-tyrosine (Fig. 1A). We introduced a bioorthogonal azide group to the C-terminus of α-tubulin by using 3-azido-tyrosine as the substrate for TTL. The desired functional molecules containing cycloalkyne were covalently conjugated by strain-promoted azide-alkyne cycloaddition (SPAAC) (Fig. 1B). To test the labeling specificity, we first conjugated Alexa Fluor 488 using this site-specific labeling (SSL) method, and separated α- and β-tubulin by high-resolution SDS-PAGE. We found that the fluorophore conjugated with the SSL method is specific to α-tubulin (Fig. 1C), as previously shown ^32^. We also found that the site-specific fluorophore labeling had little effect on the microtubule dynamic properties, even at labeling density as high as 43% (Fig. S2A, B). Using SSL-biotinylated tubulin, we further confirmed that the labeling site is indeed on the C-terminus of α-tubulin using tandem mass spectrometry (Fig. S2C). Thus, we have constructed a site-specific labeling method to covalently conjugate a functional handle to the α-tubulin CTT using recombinant TTL and commercially available reagents.

**Figure 1.**
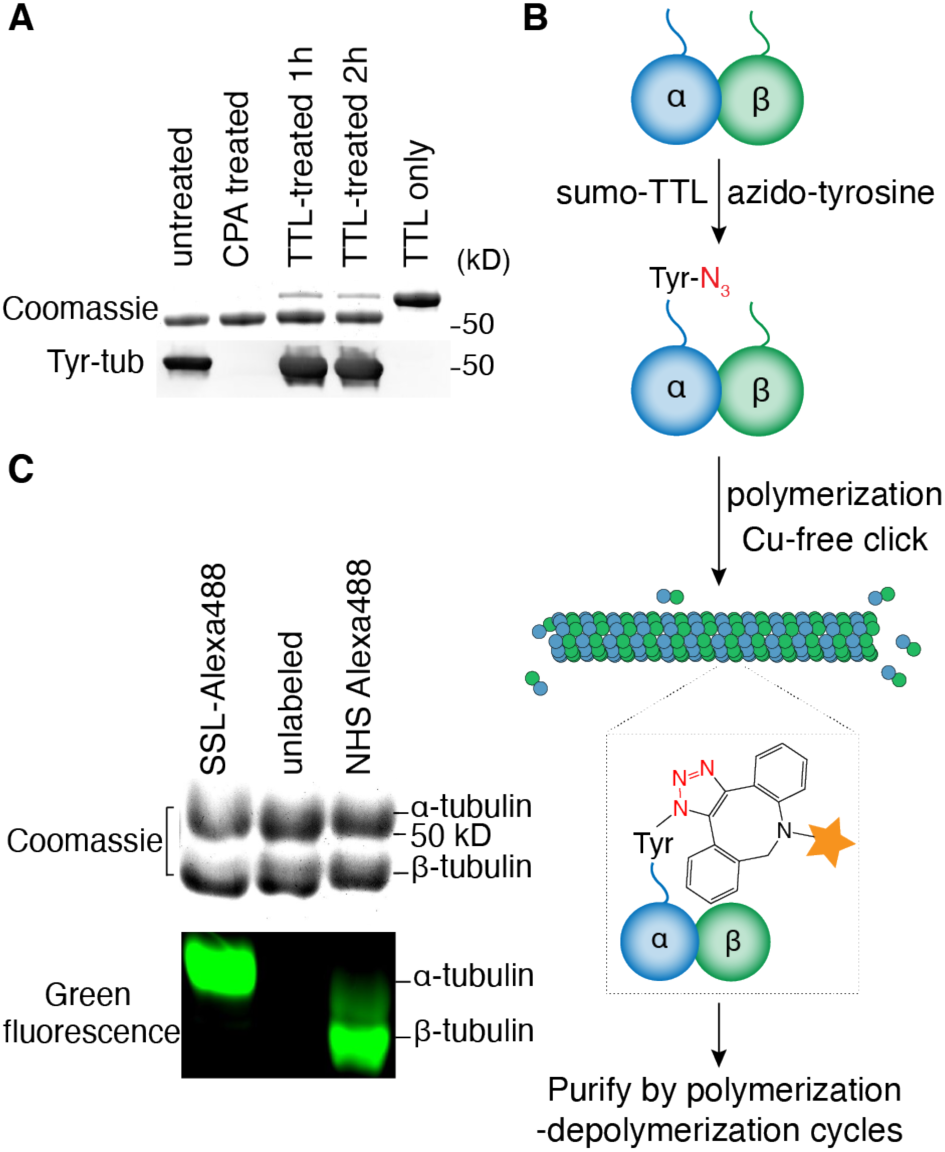
Site-specific functionalization of tubulin. (A) Recombinant tubulin-tyrosine ligase (TTL) increased the tyrosination of bovine-brain tubulin. The tyrosinated-tubulin level detected by anti-tyrosinated tubulin antibody (clone YL1/2) increased after 1 hour of incubation (bottom panel). Carboxypeptidase A (CPA) cleaves the C-terminal tyrosine of α-tubulin; CPA-treated tubulin therefore serves as a negative control. (B) Schematic process of the site-specific labeling and purification of tubulin. The orange star represents the functional moieties of interest including fluorophores, affinity tags (e.g., biotin) and macromolecules (e.g., oligo-DNAs). (C) High-resolution SDS-PAGE of Alexa-Fluor-488-labeled tubulin with either site-specific labeling (SSL) or non-specific amine-reactive labeling (labeled with NHS ester of Alexa Fluor 488). SSL-tubulin showed fluorescence signal only on α-tubulin, while tubulin labeled with NHS ester showed conjugation on both tubulin chains with a higher fluorescence signal on β-tubulin, potentially due to the higher reactivity of the β-tubulin lysine side chains.

### Pulling tubulin subunits with an optical tweezer

We next developed a tubulin-pulling assay using optical tweezers (Fig. 2). Single-stranded oligo-DNA (a GC-rich 40-mer) was covalently conjugated to the α-tubulin CTT with the aforementioned labeling method (Fig. S3A). A long, double-stranded DNA-linker (8.2 kb, 2.8-μm contour length) containing a 5’-flanking region that is complementary to the oligo-DNA handle was then hybridized to the oligo-DNA. The other end of the double-stranded DNA linkers was labeled with digoxigenin, which was bound to a polystyrene microsphere coated with anti-digoxigenin antibody. We attached the DNA linkers to taxol-stabilized microtubules, which was confirmed by total-internal-reflection-fluorescence (TIRF) microscopy and interference-reflection microscopy (IRM) (Fig. S3B). Taxol-stabilized microtubules were used because their high stability against depolymerization is compatible with the low-throughput tweezer experiments; the GDP microtubules used in the motor experiments described below are susceptible to breakage and spontaneous depolymerization, making them unsuitable for the tweezer assay. To prevent the microtubule filaments from bending or sliding when pulled by the optical trap, the microtubules were copolymerized with biotinylated tubulin. These microtubules were affixed tightly to the coverslip of a flow-channel coated with neutravidin. Anti-digoxigenin antibody-coated microspheres were then introduced to form DNA-tethers. The binding of beads to the DNA linkers was confirmed by their tethered-Brownian motions ^34^ around the microtubules.

**Figure 2.**
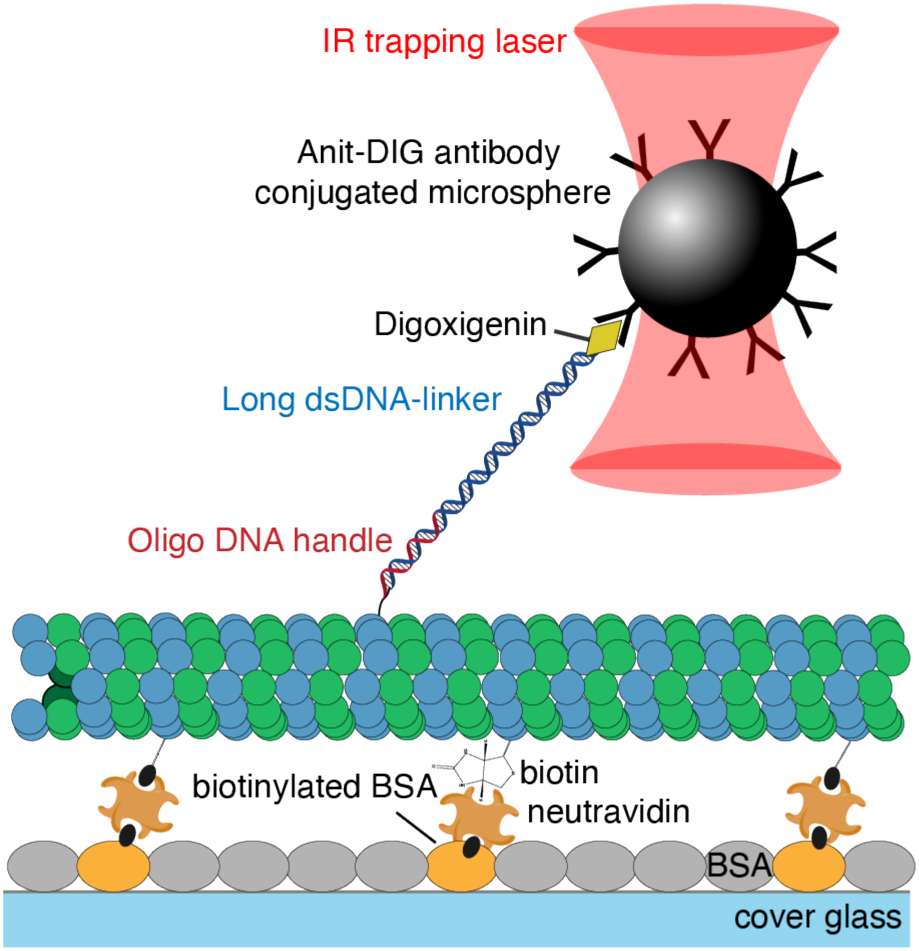
Experimental setup of the tubulin pulling assay using an optical trap. Tubulin labeled with an oligo-DNA handle using the SSL method was linked to a long DNA linker through hybridization. Tubulin conjugated to the DNA-linker was polymerized with biotinylated tubulin and affixed to the surface of neutravidin-coated coverslips. The anti-digoxigenin antibody-coated microspheres were then bound to the DNA-linker labeled with digoxigenin. Mechanical force was exerted onto the α-tubulin CTT by pulling the microsphere with the optical trap.

We pulled single tubulin subunits in the lateral direction parallel to the coverslip surface and perpendicular to the microtubule axis using an optical trap. We chose this pulling direction mainly to avoid the attachment of more than one DNA linker to the bead during the pulling experiment. An example of a single-molecule force-extension trace is shown in Fig. 3A (median-filtered trace in orange). The low-force region of the force-extension curves corresponds to the stretching of the long double-stranded DNA linkers and was well described by the worm-like chain model, which included an elasticity term (Fig. 3A, green curve) ^35,36^. We observed both stepwise lengthening and ruptures in the pulling traces (Fig. 3A, arrows). After the rupture events, the polystyrene microsphere diffused away after the laser trap was turned off, showing the bead was no longer tethered to the microtubule. Stepwise-lengthening events were often observed prior to the rupture (Fig. 3A, 44 out of 48 traces with rupture events). The lengthening events were absent in control force-extension traces in which DNA-linkers were tethered directly to the surface (zero out of 37 traces; see example of a control DNA-linker pulling trace in Fig. S4). We therefore interpreted stepwise lengthening as partial unfolding of tubulin subunits.

**Figure 3.**
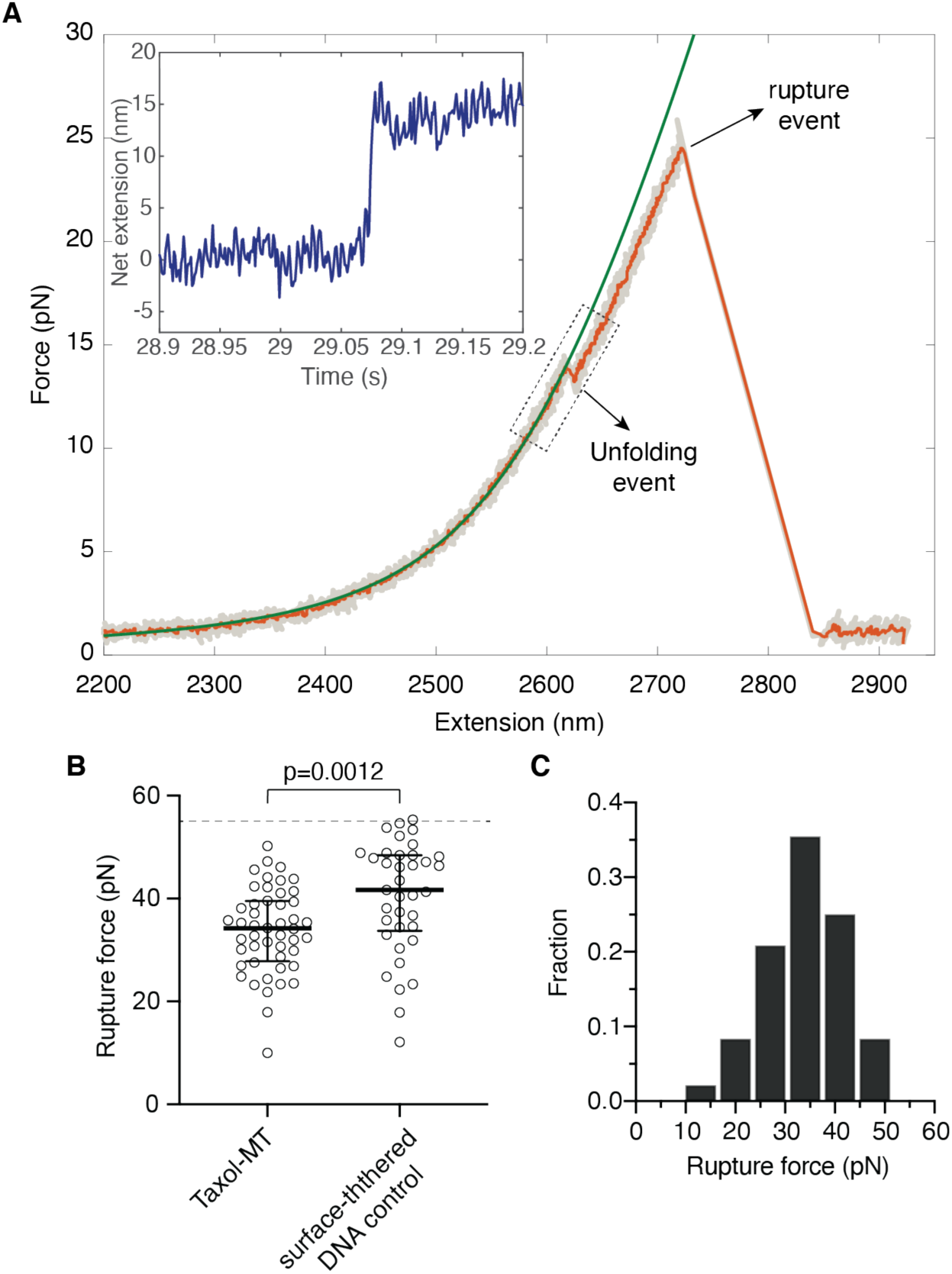
Pulling tubulin subunits from the taxol-stabilized microtubule lattice with an optical trap. (A) Example of a force-extension curve from the tubulin-pulling assay. Unfolding and rupture events are indicated by arrows. Experiments were performed with a constant pulling velocity of 0.32 μm/s, using an 8.2 kb-DNA-linker (contour length 2.8 μm). Gray: raw trace; orange: median-filtered trace; green: worm-like chain model fit up to the unfolding event. Inset: the extension after subtracting the extension of DNA linker showed a clear unfolding step of ∼15 nm. The corresponding region of the force-extension curve is marked by a rectangle. (B) Rupture forces from the taxol-MT pulling traces were significantly lower than the ones from the surface-tethered DNA linker controls (*p* = 0.0012, Mann-Whitney *U* test). Note that the average rupture forces measured from the linker-only control were likely to be an underestimation because the measurements were capped by the maximum force that could be generated by our optical trap (∼55 pN; dashed line in Fig. 3B). (C) Rupture-force histogram in the taxol-stabilized microtubule pulling experiments.

The rupture events occurring at forces 33.8 ± 1.2 pN, (mean ± SE unless otherwise noted, *n* = 48 traces; Fig. 3B points to the left) are primarily due to removing tubulin from the microtubule lattice, based on the following arguments. First, breaking the covalent bond between the oligonucleotide and the tubulin protein is expected to require much higher forces (on the nanonewton scale ^37^). Second, the shear force to break the 37-nucleotide oligoDNA-DNA linker interaction is expected to be ∼60 pN ^38^, much higher than the measured rupture forces. And third, the force to break the antibody-digoxigenin bond must be larger than the force required to rupture the surface-tethered DNA bead controls, which also used the same anti-digoxigenin antibody (Fig. 3B points to the right; 40.3 ± 1.8 pN; *n* = 37 traces). Thus, a great portion of the rupture events observed in the taxol-microtubule pulling experiments likely correspond to the removal of tubulin by the applied force.

From the distribution of rupture forces (Fig. 3C), we estimated that the force typically required to remove tubulin from the taxol-stabilized microtubule lattice to be around 30 pN at this pulling velocity (∼0.32 μm/s). A complication of our optical trap assay is the uncertainty of the actual pulling direction with respect to the tubulin being pulled as we were unable to identify the protofilament on which the subunit was located. The force required to extract tubulin when pulling in the optimal direction (perhaps orthogonal to the lattice) is therefore likely to be lower than the rupture forces measured in our optical tweezer assay.

### Unfolding of tubulin subunit under mechanical force

To estimate the unfolding step size, we subtracted the length of the DNA linker estimated by fitting the worm-like chain model (Fig. 3A, green curve). The net extension showed clear stepwise increases, which we interpret as the partial unfolding of tubulin (Fig. 3A, inset). The typical step size was 10-20 nm (3% to 92% range) (Fig. 4A), equivalent to the unfolding of ∼25-50 amino acids of the polypeptide (∼0.4 nm/residue, ^39^). Examining the secondary structure of α-tubulin close to the C-terminus ^40^, we hypothesize that these events corresponded to the unfolding of the last C-terminal 2 to 3 helices of α-tubulin (H11, H11’, H12, blue in Fig. 4B); this is consistent with an earlier coarse-grained molecular simulation of α-tubulin pulling trajectories ^41^.

**Figure 4.**
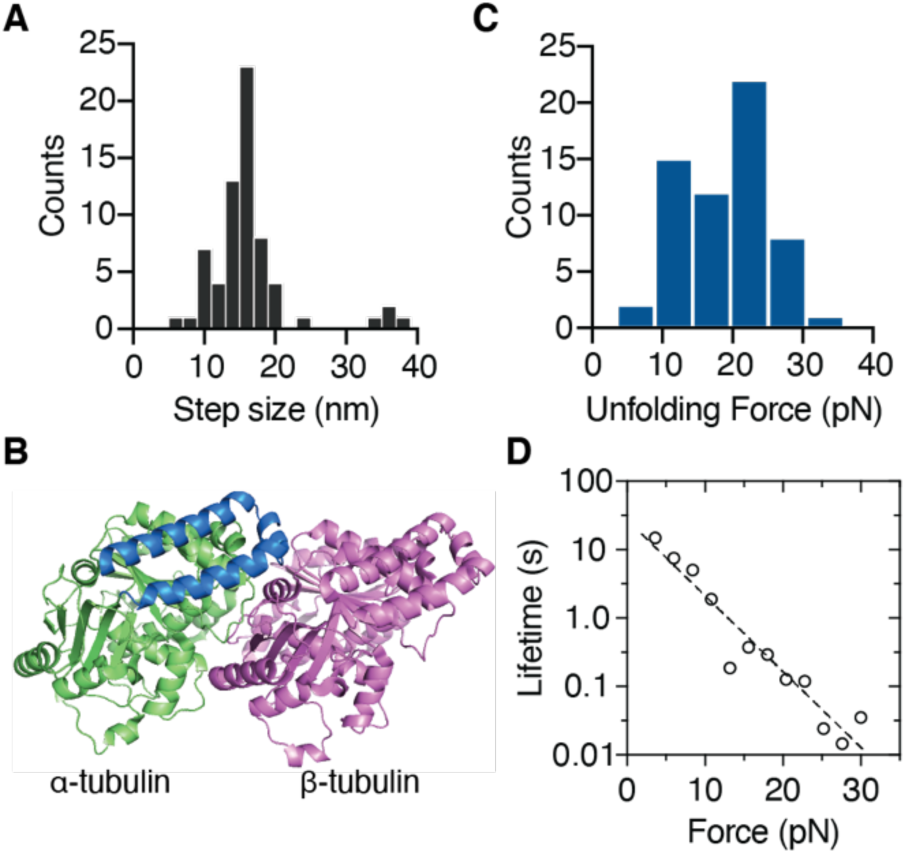
Characterization of the unfolding events of tubulin during the microtubule-pulling experiments. (A) The distribution of the unfolding step sizes (16 ± 6 nm; mean ± SD, *n* = 66 unfolding events). (B) Structure of αβ-tubulin looking at the microtubule surface (PDB: 3J6F, ^44^). The hypothesized unfolding 50 amino acids of α-tubulin is highlighted in blue (Helix 11, 11’, 12). (C) Distribution of unfolding forces associated with step sizes between 10 to 20 nm (19.2 ± 0.8 pN; mean ± SE, *n* = 60 events). (D) Lifetime of the folded state *τ* as a function of force *F* transformed from the unfolding force histogram in (C) (circles). Fit with the Bell equation (dashed line): *τ*_0_ = 25.9 ± 1.6 s, *x*^‡^ = 1.05 ± 0.10 nm (SE); *R*^2^ = 0.92.

To obtain more detailed kinetic information about the unfolding events, we transformed the unfolding forces associated with the 10-20 nm step sizes (Fig. 4C) into the force-dependent lifetime of the folded state τ(*F*) (Fig. 4D) using the method of ^42^. Though the pulling axis in our optical trap assay is likely to be a poor reaction coordinate, the force-dependent lifetime was nevertheless well-described by Bell’s model 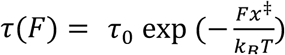 ^43^, with the lifetime of the folded state at zero force *τ*_0_ = 25.9 ± 1.6 (s) and the distance to the transition state *x*^‡^ = 1.05 ± 0.10 nm (Fig. 4D). Thus, these results suggest that the unfolding of the region spanning H11 to H12 of α-tubulin is an intermediate step in the tubulin-removal pathway, and the spontaneous unfolding of this region takes place on the timescale of tens of seconds.

### Tubulin extraction from GDP-microtubules with molecular motors

Because the mean force required to extract tubulin with the optical tweezers (34 pN) is not so different from the force to rupture the DNA-bead tether (41 pN), we sought an alternative method to measure the tubulin-removal force. In addition, we sought a high-throughput assay, unlike the tweezer assay in which we can only pull on one tubulin at a time. We therefore developed a motor-pulling assay in which many force-events can be measured simultaneously ^45^. To this end, we first prepared GDP-microtubules capped with GMP-CPP tubulin (GMP-CPP is a slowly hydrolysable GTP analogue) to prevent their spontaneous depolymerization. The GDP-microtubules were prepared by copolymerizing tubulin conjugated to biotinylated DNA linkers (3.8 kb) along with digoxigenin-labeled tubulin in the presence of GTP, which subsequently hydrolyzes to GDP (see Materials and Methods for details). These microtubules were bound to the surface by anti-digoxigenin antibody (Fig. 5A). Biotinylated kinesin-1 (a truncated rat kif5C) was then joined to the DNA linkers via neutravidin (Fig. 5A). Note that depending on the concentration, the number of kinesin-1 motors per DNA linker varies from 1 to 3 (there are four biotin-binding sites per neutravidin tetramer with one used to attach the DNA linker). We stained the DNA linkers with the fluorescent DNA intercalator SYTOX Green and visualized it using TIRF microscopy. In the presence of ATP, the DNA linkers, which formed a compact random coil of diameter ∼0.5 μm, were quickly stretched along the microtubule filament by the kinesins (Fig. 5B,C). The fluorescence intensity of the DNA linker increased as it approached its contour length due to the tension-dependent enhancement of affinity to SYTOX green ^46,47^. Thus, the stretching of the tubulin-DNA complexes by the motors could be directly visualized.

**Figure 5.**
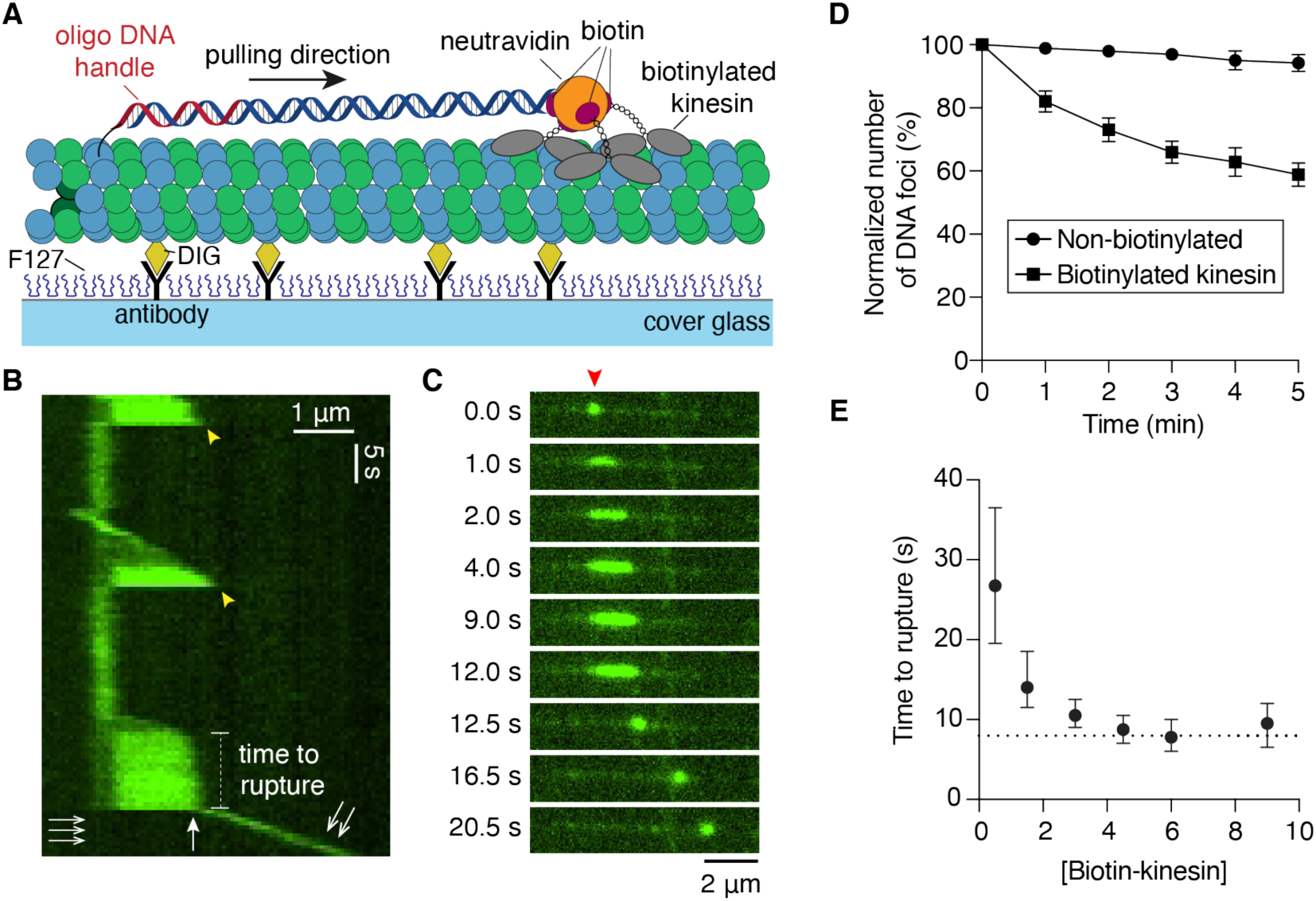
Kinesin-1 pulled out tubulin subunits from the GDP-microtubule lattice. (A) Experimental scheme of the motor-pulling assay. GDP-tubulin subunit attached to a DNA linker (3.8 kb) was pulled by biotinylated kinesin-1 molecules along the microtubule axis. The GDP-microtubules were stabilized with GMPCPP-tubulin caps on both ends (not shown for simplicity). (B) Example kymograph of kinesin-1 stretching a DNA-linker (stained by SYTOX Green) and pulling out a GDP-tubulin subunit. The dissociation event of kinesin-1 is indicated with yellow arrowheads. The rupture event corresponding to the dissociation of tubulin was marked by the white arrow. (C) Time-lapse images of the last stretching round before rupture from the kymograph in (B). The DNA molecule of interest is indicated with a red arrowhead. (D) The number of microtubule-linked DNA molecules remaining after introducing 6 nM of kinesin-1 (either non-biotinylated or biotinylated with ∼50% stoichiometry) imaged at 1 frame per minute (error bars: SDs). Biotinylated kinesin-1 led to a significant reduction of fluorescent DNA molecules after 5 minutes as compared to the non-biotinylated kinesin-1 control (Welch *t*-test *p* = 0.0003; percentage of DNA remained: 59 ± 4% vs. 94 ± 3%; mean ± SD, *n* = 3 experiments). (E) Time-to-rupture (median ± 95% confidence interval) decreased with increasing concentration of biotinylated kinesin-1 to a value of ∼8 s (dashed line). Data collected from triplicate experiments with *n* = 108, 105, 130, 122, 112, 99 events at each concentration (from 0.5 nM to 9 nM).

After the DNA linker was fully stretched, the motors stalled for a variable time until one of three events occurred. (i) The stretched DNA recoiled to the anchor point and remained attached to the microtubule (yellow arrowheads in Fig. 5B), presumably because the motors dissociated from the microtubule lattice. (ii) The stretched DNA recoiled and was transported along the microtubule filament by the kinesins (double arrow in Fig. 5B), presumably due to the removal of the GDP-tubulin subunit from the lattice (Fig. 5B white arrow, 5C). Note that no DNA remained associated with the anchor point (triple arrow in Fig. 5B). (iii) The stretched DNA separated into two parts, one part recoiling at the anchor point on the microtubule and the other part recoiling and moving along the microtubule (see an example in Fig. S5A). This third event we attribute to photocleavage of the DNA. As further evidence that the motors are actively pulling the entire linker DNA off the microtubule lattice (presumably with the tubulin attached to it), we quantified the number of DNA molecules remaining on the microtubules over time by imaging at low frequency (snapshots with 1 minute interval) to minimize photobleaching. After introducing biotinylated kinesin-1 and ATP, the number of DNA molecules conjugated to the microtubules decreased by ∼40% in 5 minutes, while more than 90% of the DNAs remained when the same concentration of non-biotinylated kinesin-1 (and ATP) was used (Fig. 5D). This decrease was not a result of tension-enhanced photocleavage because most of the DNA molecules were in the coiled state. This result demonstrates that DNA is removed by biotinylated motors and not by photocleavage.

We measured the time-to-rupture at different concentrations of biotinylated kinesin. The median rupture time decreased with increasing kinesin concentration and approach a plateau of ∼8 s (Fig. 5E). Consistent with this, the rupture-force histogram at saturating concentrations of biotinylated kinesin (4.5 to 9 nM) was well fit by an exponential with a decay time of 8.8 s (Fig. S6A). The long attachment time of 8 s is consistent with kinesin having a catch-like association with the microtubule in which the lifetime of the pulling state increases with load along the axis of the microtubule ^48,49^.

To compare the optical-tweezer and motor assays, we transformed the rupture-force histogram measured in the optical tweezer assay (Fig. 3C) into a force-dependent bond lifetime using the method of ^42^: 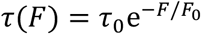, where *τ*_0_ = 37 s is the breakage time in the absence of force and *F*_0_ = 6 pN is the force that decreases the lifetime e-fold. The 8-s rupture time measured in the motor assay corresponds to a force of 8.5 pN (Fig. S6B). This is equivalent to the maximum force generated by 1-2 kinesins (one kinesin generates 4 to 7 pN, ^50^), assuming that the kinesin force is proportional to the number of motors ^49^. Given that we observed stepwise increases of fluorescence even at the highest motor densities (Figure S5B, C), which we interpret as increasing numbers of motors, not all three kinesins are bound to the microtubule and generating force during the tubulin extraction process. Therefore, 1 to 2 kinesins generating an 8.5-pN force that pulls tubulin out of the lattice in ∼8 s is reasonable. Thus, we believe there is good qualitative agreement between the tweezer and motor assays, despite the technical limitations of our optical tweezer assay (e.g., some ruptures may be due to breaking the antibody-digoxigenin bond, and the reaction pathway for tubulin extraction may be more complex than just a single energy barrier) and the details of the experiments differ (e.g., GDP- vs taxol-stabilized microtubules, and different force vectors relative to the microtubule axis). In conclusion, our results suggest that the forces generated by 1 to 3 kinesin-1s are able to extract GDP-tubulin from the lattice in several seconds.

## Discussion

Using single-molecule techniques, we have shown that tens of piconewtons of mechanical force can partially unfold and remove tubulin subunits from the microtubule lattice on the second timescale. The distribution of rupture forces associated with tubulin removal measured by optical tweezers is consistent with Bell’s equation in which the lifetime of ∼40 s at zero force decreases e-fold per 6 pN as the force increases. The lifetimes measured in the high-throughput motor assays in which one to three kinesins pull on tubulin are consistent with the optical-tweezer measurements. We now evaluate the implications of these tubulin-extractions forces on the stability of microtubules and the mechanism of microtubule severing proteins such as spastin and katanin.

Structural studies suggest that AAA unfoldases, such as ClpXP, ClpB, Vps4 and spastin/katanin, translocate 2 amino acids (∼0.8 nm) through their central pores per ATP hydrolyzed ^20,22,24^. If these enzymes are well coupled under load, they could potentially generate forces as high as 120 pN (Δ*G*_ATP_/0.8 nm), sufficient to rapidly unfold and remove tubulin from the microtubule lattice. Indeed, ClpXP ^18,19^ and ClpB ^17^ can generate forces up to 20 and 50 pN respectively, sufficient to partially unfold and extract tubulin. While the force generated by the microtubule severases remains unknown, our force spectroscopy of tubulin supports the plausibility of spastin and katanin using an unfoldase-type mechanism where mechanical force exerted on the tubulin C-terminal tails threads tubulin through the pore, partially unfolding and then pulling tubulin subunits from the lattice ^12,51^. Whether severing enzymes thread the entire tubulin polypeptide chain through their pore is less clear: with an ATP hydrolysis rate of 22 s^-1 52^, threading the full chain would take spastin about 10 s.

The motor pulling assay demonstrated that the forces generated by several kinesin-1 molecules are sufficient to extract GDP-tubulin from the lattice. Removal of tubulin from the unstabilized GDP-tubulin lattice did not lead to immediate microtubule depolymerization (within minutes), however. This suggests that the lattice destruction observed when dyneins and kinesin-14s walk along microtubules, and when microtubules glide across surfaces coated with these motors must be due to accumulated damage to the lattice by several motors ^26^. For cellular processes such as mitosis ^53^ and tissue development ^54^ that involve frequent sliding of microtubule filaments by multiple motors, the removal of tubulin under force may create defects on the microtubules and require the reincorporation of new GTP-tubulin to the lattice to prevent filament breakage ^30^.

How frequently will a tubulin extraction event occur when a single molecular motor walks on the microtubule track? We expect a tubulin extraction event to take at least 30 s when the stall force (4 to 7 pN) of a single kinesin is applied (based on the rupture time measured in the presence of lower concentration of biotinylated kinesin-1, Fig. 5E, 0.5 nM). The time could be much longer if the force spectroscopy probes only the first step in the extraction pathway, and this step is reversible in the absence of force. The presence of more than one reversible step prior to tubulin removal likely explains why the tubulin subunits in a microtubule do not undergo a complete turnover within a minute in the absence of force as suggested from the Bell’s model fit (Fig. S6B, lifetime at zero force ∼40 s). For motors walking without load, the force is expected to be ∼1 pN, a lot less than the stall force ^55^, and so the lifetime will be even longer (>1 minute). Given that the kinesin run-time along the microtubule is ∼1 s, we expect that only about 1% of kinesin runs will result in tubulin removal. Therefore, a large number of the GDP-microtubules are expected to survive for several minutes before they depolymerize even in the presence of high motor concentrations ^26^.

In conclusion, our results provide the first experimental investigation of the mechanical force required to unfold and extract tubulin from the microtubule lattice. Several important questions remain to be explored. For instance, how many tubulin dimers are removed from the microtubule lattice – just one or multiple? Does the partial unfolding of tubulin weaken tubulin-tubulin bonds, facilitating the dissociation of subunits? How does the unfolding and rupture force depend on the pulling orientation? Is there a difference in force requirement when pulled on the C-terminal tail of β-tubulin as compared to α-tubulin? We expect our assay design to provide an important step toward understanding the molecular response of tubulin to mechanical force, with important implications in the mechanics of the microtubule cytoskeleton and its cellular functions.

## Materials and Methods

### Protein preparation and assays

Bovine brain tubulin was purified in house based on the method described previously ^56^. Codon-optimized human tubulin-tyrosine ligase (TTL) was custom synthesized (IDT) and cloned into a pET vector with N-terminal His_6_-SUMO tag (Addgene plasmid #29659; a gift from Scott Gardia, University of California Berkeley) by ligation-independent cloning. His_6_-SUMO-TTL was expressed in *Escherichia coli* (BL21-DE3 competent cells, Agilent) overnight at 18 °C in LB broth (induced by 0.5 mM IPTG). The cells were harvested and stored at -80 °C until purification. To purify His_6_-SUMO-TTL, the cells were resuspended in cold lysis buffer (20 mM NaH_2_PO4, 0.2 M NaCl, 5 mM imidazole, pH 6.9, 0.2 mM pefabloc, 0.1 mg/mL lysozyme, 1 mM dithiothreitol (DTT), 0.3 U/μL benzonase), and lysed by sonication on ice. The lysate was clarified by centrifugation and loaded onto a HisTrap column (GE Healthcare). The column was washed with the imidazole buffer (20 mM NaH_2_PO4, 0.2 M NaCl, 20 mM imidazole, pH 6.9, 1 mM DTT). The protein was then eluted with a continuous gradient (20 mM to 500 mM imidazole). The eluted protein was concentrated by centrifugal filters (30 kD cutoff) and further purified by size exclusion chromatography (SEC). The protein was eluted from size exclusion column with SEC buffer (20 mM MES-KOH, 0.1 M KCl, 10 mM MgCl_2_, 1 mM EGTA, 1 mM DTT, pH 6.9. The purified protein was concentrated with centrifugal filters, aliquoted, flash frozen with liquid nitrogen and stored at -80 °C.

To test the tyrosination activity of recombinant TTL, 1 μM of purified His_6_-SUMO-TTL was incubated with 10 μM of bovine tubulin in the presence of 2.5 mM ATP, 1 mM L-tyrosine and 5 mM DTT in BRB80 (80 mM PIPES-KOH, 1 mM EGTA, 1 mM MgCl_2_, pH 6.9) at 37 °C. A negative control without tyrosinated tubulin was prepared by treating 3 mg/mL bovine tubulin with 0.02 mg/mL carboxypeptidase A (CPA, Sigma Aldrich) on ice for 15 min to proteolytically cleaved the C-terminal tyrosine of α-tubulin. The tyrosinated tubulin level was detected by western blot using monoclonal anti-tyrosinated tubulin antibody (clone YL1/2; 1:2500 dilution; EMD Millipore) as primary antibody and a goat anti-rat secondary antibody (alkaline phosphatase-conjugated, 1:1500; Invitrogen).

For the motor-pulling assay, we used a truncated rat kinesin-1 (the first 430 aa of rat Kif5C, here denoted as rk430) ^57^ fused with a mScarlet-SNAP-His_6_ tag at the C-terminus. The expression of rk430-mScarlet-SNAP-His_6_ was performed in the same way as TTL as described earlier. The cell pellet was resuspended in cold lysis buffer (30 mM HEPES-NaOH pH 7.4, 10 mM imidazole, 0.3 M NaCl, 1 mM DTT, 10 μM ATP) containing protease inhibitor (Roche) and 0.3 mg/mL lysozyme. The cells were then lysed by sonication and clarified as earlier described. The clarified lysate was flowed through a HisTrap column and eluted with a gradient of imidazole (from 10 to 500 mM). The peak fractions were collected, desalted with Zeba desalting column (Thermofisher) into anion-exchange buffer (30 mM HEPES-NaOH pH 7.4, 1 mM DTT, 10 μM ATP) and bound to a HiTrapQ column (GE Healthcare). The proteins were eluted with a NaCl gradient (0.01 to 1 M). The peak fractions were collected, concentrated using an Amicon centrifugal filter and purified further with SEC, eluting with BRB80 containing 10 μM ATP and 0.1% β-mercaptoethanol. The purified protein was flash frozen with liquid nitrogen as small aliquots and stored at -80 °C. The activity of purified kinesin was confirmed by the single-molecule fluorescence stepping assay as previously described ^58^.

### Site-specific labeling by tubulin-tyrosine ligase (TTL)

To introduce the bio-orthogonal azide functional group, bovine brain tubulin was mixed with His_6_-SUMO-TTL (5:1 molar ratio of tubulin: TTL) in the presence of 1 mM 3-azido-tyrosine (Watanabe Chemical), 2.5 mM MgATP in BRB80 and incubated at 37 °C for 45 to 60 minutes. To remove TTL and excessed azido-tyrosine, the labeled tubulin was polymerized into microtubules by adding GTP (final concentration 1 mM), glycerol (final concentration 33.3%) and MgCl_2_ (final concentration 3.5 mM) at 37 °C for 30 min. The microtubule solution was layered on a glycerol cushion (BRB80 with 4 mM MgCl_2_, 1 mM GTP, 60% glycerol) and centrifuged (340,000 x g, 35 min at 35 °C). The pellet was resuspended in warm click-labeling buffer (BRB80, 1 mM GTP, 4 mM MgCl_2_, 40% glycerol), followed by the addition of oligo-DNA or small molecules conjugated to dibenzocyclootyne (DBCO) (∼0.5 mM final concentration for organic fluorophores (ThermoFisher) or biotin with PEG_4_ linker (Sigma Aldrich); or ∼0.15 mM oligo DNA: 5’-TGGACTGATGCGGTATCTGCGATATCCTACGCAGGCGTTT-3’-DBCO (synthesized from IDT). The copper-free click reaction took place at 37 °C for 1 hour with gentle shake and occasional mixing. The labeled microtubules were then again layered on the 60% glycerol cushion and centrifuged at 35 °C (446,000 x g, 20 min) to remove the unreacted labeling reagent. The microtubules were resuspended with BRB80 and incubated on ice for 30 min to induce depolymerization. The solution was centrifuged at 4 °C (285,500 x g, 10 min) to remove aggregates or precipitates. An additional polymerization and depolymerization cycle was used to further purify functional tubulin based on a previous protocol ^59^. The labeled tubulin was flash frozen in liquid nitrogen and stored at -80 °C. The concentration of tubulin and labeling density of fluorophores was determined by UV-Vis absorption. Biotinylation level was quantified using a biotin quantification kit (Thermofisher). The oligo-DNA labeling density was estimated by measuring the band intensities from SDS-PAGE. Note that we chose copper-free click chemistry due to the commercial availability of azido-tyrosine and its faster reaction kinetic without the need of catalyst as compared to the hydrazone formation strategy used in ^32^.

To confirm the labeling specificity, high-resolution SDS-PAGE was used to separate α-and β-tubulin as previously described ^60^. To examine the specific labeling site, SSL-biotinylated tubulin was first buffer exchanged into phosphate buffer saline (PBS) pH 7.4. Biotinylated tubulin was then protease digested by AspN (Promega) at 37 °C overnight (1:50 w/w ratio). An equal volume of BRB80 was added to stop the protease digestion, and incubated with NeutrAvidin agarose resin (Pierce) for 1 hour at room temperature to pull down the biotinylated peptides. The resin was washed with PBS and water, and the biotinylated peptides were eluted by incubating the resin with an aqueous solution containing 80% acetonitrile, 0.2% trifluoroacetic acid, 0.1% formic acid at 60 °C for 5 min. The elute was then sent for proteomic analysis. The peptides were separated using reverse-phase ultra-high pressure liquid chromatography (UPLC) and analyzed by tandem mass spectrometry (MS/MS) with electrospray ionization. MS/MS spectra were searched by MASCOT against the bovine α-tubulin sequence with the modifications corresponding to the copper-free click biotin adduct on tyrosine and polyglutamylation (up to 4 glutamate chains) on glutamate side chains. All biotinylated peptide hits corresponded to the C-terminal tail of α-tubulin, confirming the site-specificity of the labeling method. Microtubule dynamic assays were used to measure the dynamic properties of SSL-Alexa Fluor 488-tubulin and unlabeled tubulin (in BRB80 supplemented with 1 mM GTP and 5 mM DTT) as described previously ^61,62^ using interference-reflection microscopy (IRM) ^63^.

### Microscopy setups

The two-color (488 nm and 561 nm excitation lasers) total-internal-reflection-fluorescence (TIRF) microscopy and interference-reflection microscopy (IRM) was set up on a Nikon Ti-Eclipse microscope as previously described ^63^ with a sCMOS camera (Zyla 4.2 Plus, Andor) for both TIRF and IRM imaging.

The home-built optical tweezer setup was constructed around a Zeiss inverted microscope as previously described ^64^ with the near infrared 1064 nm Nd:YAG laser (IPG photonics). The system was coupled to a differential-interference-contrast (DIC) condenser to visualize microtubule filaments and the polystyrene microspheres using blue light LED illumination ^65^. We used a Zeiss Plan-Neofluar 100x/1.3 NA objective for both trapping and DIC imaging. The sample stage was controlled by a three-axis piezoelectric stage with sub-nanometer precision (Physik Instrumente). To detect the position of the bead with nanometer sensitivity, we used a quadrant photodiode (QPD; First Sensor) for back-focal-plane detection and calibrated the position detection and trap stiffness using a combined drag force-power spectral analysis method as described in ^64,66^. All optical tweezer experiments were performed at similar trap stiffness (∼0.28-0.32 pN/nm) and the time traces were recorded at 1 kHz sampling frequency. The optical tweezer system was operated by custom-written software using LabView (National Instruments) ^64^.

### Preparation of microspheres

The carboxylated polystyrene microspheres (0.58 μm diameter; Bangs Laboratory) were labeled with anti-digoxigenin Fab fragment (Roche) based on a two-step functionalization method described in ^67^ with the following modifications. First, we used a 3:1 molar ratio of 2 kDa α-methoxy-ω-amino PEG: 3 kDa α-amino-ω-carboxy PEG (Rapp Polymere) in the first functionalization step. Second, we used 0.16 mg/mL anti-digoxigenin Fab in phosphate buffer saline (PBS) pH 7.4 for the antibody conjugation step. The Fab-conjugated microspheres were sonicated briefly (∼15 sec) and incubated with 1 mg/mL BSA on ice for at least 10 min prior to the optical tweezer experiments to enhance the bead surface passivation.

### Preparation of DNA linkers

The long DNA linkers containing a 5’ overhang on one end and digoxigenin on the other end were prepared by PCR ^68,69^ using Q5 polymerase (NEB). Lambda phage DNA (Thermofisher) was used as the PCR template. The digoxigenin-labeled primer was prepared by labeling 0.25 mM oligo containing 3 amino groups (amino groups at the 5’ end and the underlined bases: TC<u>T</u>AAG<u>T</u>GACGGCTGCATACTAACC; synthesized by IDT) with 66 mM digoxigenin N-hydroxysuccinimide ester (Roche) in 50 mM HEPES-NaOH, 67% DMSO, pH 8.3 overnight with shaking at room temperature. The labeled primer was purified by G-25 microspin column (Cytvia) and Monarch DNA purification kit (NEB). The successful conjugation of three digoxigenin moieties was confirmed by the shift of the molecular weight using denaturing urea polyacrylamide gel electrophoresis. The primer with 5’-overhang was custom synthesized (IDT) with the following sequence: <u>CGCCTGCGTAGGATATCGCAGATACCGCATCAGTCCA</u>XCAACGGTCGATTGCCTG ACGGA where the sequence complementary to the oligo handle labeled on tubulin is underlined and X is the abasic residue (1’,2’-dideoxyribose). This primer pair produced the 8.2 kb DNA linker used for the optical tweezer taxol-stabilized microtubule pulling assay. For surface-tethered DNA control, the purified 8.2 kb DNA linker was annealed to the oligo DNA handle with 3’-biotin (5’-TGGACTGATGCGGTATCTGCGATATCCTACGCAGGCGTTT-3’-biotin) in PBS at room temperature for 3 hr followed by purification using Monarch DNA purification kit (NEB).

For the motor pulling assay, the digoxigenin-labeled primer was replaced with 5’-biotinylated primer (IDT). The complete primer pairs and the corresponding product length is provided in Table S1. The DNA linkers produced by the PCR reaction were first desalted using an amicon centrifugal filter and purified using the PureLink PCR purification kit (Thermofisher). DNA linkers were eluted with low TE buffer (10 mM Tris-HCl, 0.1 mM EDTA, pH 8.0) and stored at -20 °C as small aliquots.

### Taxol-stabilized microtubule pulling assay with optical tweezer

To prepare taxol-stabilized microtubules conjugated to DNA linkers, the tubulin-DNA mixture with 20 μM of tubulin labeled with oligo DNA (labeling density ∼3%), 4 μM of biotinylated tubulin (labeling density ∼50%), ∼3.4 nM 8.2-kb DNA linker was incubated on ice for 15 min and polymerized at 37 °C in the presence of 4 mM MgCl_2_, 2% DMSO, 1 mM GTP for 25 min. The polymerized DNA-microtubules were quickly diluted into BRB80 supplemented with 10 μM taxol and pelleted by centrifugation. DNA-microtubules were then resuspended in BRB80 containing 10 μM taxol.

The flow channel was prepared by using one 18×18 and one 22×22 mm^2^ Piranha solution-cleaned and silanized coverslips sandwiching parafilm stripes as previously described ^62,70^. The flow channel was incubated with 0.02 mg/mL biotinylated bovine serum albumin (BSA) (Sigma Aldrich) solution for 5 min and washed by BRB80. The channel was then passivated by 3% BSA solution for 30 min and washed with BRB80+1%Tween20. The chamber was further incubated with 0.05 mg/mL Neutravidin solution containing 1 mg/mL BSA and 1% Tween20 for 10 min and washed again with BRB80+1%Tween20. Taxol-stabilized DNA-microtubules were then attached to the surface under solution flow. Polystyrene beads conjugated with anti-digoxigenin Fab (∼0.01% beads in BRB80 supplemented with 0.1%Tween20, 1 mg/mL BSA and 10 μM taxol) were then introduced to the chamber and incubated for 30 min at room temperature to allow the attachment of DNA linkers. The unbound beads were then washed out. Freshly prepared oxygen scavenger solution [40 mM glucose, 0.04 mg/mL glucose oxidase (from *Aspergillus niger*; Sigma Aldrich), 0.02 mg/mL catalase (from *Aspergillus niger;* EMD Millipore), 0.2 mg/mL casein, 10 mM DTT, 10 μM taxol, 0.1% Tween 20] was then introduced into the channel. The channel was then sealed with VALAP (equal ratio of Vaseline, lanolin and parafilm) for the optical tweezer experiments. Tethered beads undergoing tethered Brownian motion close to the microtubules (imaged by DIC) were selected for optical trapping and the calibration was performed for each bead. The tethered bead was trapped at ∼300 nm away from the surface and the stage was first set to oscillate with a small amplitude (2.5 to 2.8 μm) in the lateral direction of the microtubule at 0.1 to 0.2 Hz. The position of the stage was finely adjusted so that the trap center was aligned with the anchor point of the linker based on the symmetry of the QPD voltage-distance traces. The amplitude we used in this process was quite small so that the force was typically less than 5 pN. To collect the pulling traces, the stage was oscillating with constant velocity (∼0.32 μm/min) with an amplitude of 3.2 or 3.3 μm until rupture was observed (typically within 1 or 2 cycles). More than 95% of the traces showed rupture events in our experiments. All traces were collected within 60-75 min after the oxygen scavenger solution was introduced.

For the surface-tethered DNA control experiments, 22×22 mm^2^ coverslips were first cleaned by three cycles of sonication in 1 M KOH and ethanol (15 min each) and dried under nitrogen gas. To covalently functionalize the cleaned coverslips, 40 μL of biotinylation solution [1 mg/mL of biotin-PEG5000 silane, 100 mg/mL PEG5000-silane (Nanocs) dissolved in 95% ethanol] was sandwiched by two coverslips and incubated at room temperature for 2 hr inside a sealed wet chamber. The functionalized coverslips were then rinsed with water, dried under nitrogen gas and stored. The flow channel was constructed in the same method as the taxol-microtubule pulling experiments. The channel was passivated with 3% BSA and washed with BRB80+1%Tween20. To tether the biotinylated DNA-linker, 10 μg/mL Neutravidin solution containing 1 mg/mL BSA and 1% Tween 20 was then introduced to the channel followed by another wash of BRB80+1% Tween 20. The channel was then incubated with 15 pM of biotinylated DNA linkers for 15 min followed by another wash. The successful hybridization of the DNA linkers was verified by staining of dsDNA (QuantiFluor, Promega). Bead tethering and optical tweezer pulling were performed as the taxol-stabilized microtubule pulling assay described above.

### Analysis of pulling traces

To obtain the force-extension curves from the pulling traces, we employed the strategy described in ^35^ to identify the anchor point based on the symmetry of pulling traces during stage oscillation. The extensions were calculated by taking the pulling geometry into account. Due to the long contour length of the DNA linker, the total force was within ∼10% difference from the force along the pulling direction. An 11-point moving median filter was applied to the force-extension traces and the traces were fit with the worm-like chain (WLC) model ^35,36^(up to the unfolding steps if they were present). Traces that showed evident asymmetry during the stage oscillation cycles or deviated significantly from the predicted DNA force-extension curve were discarded. Rupture forces and unfolding forces were measured from the sharp decrease of the force from the force-extension curves and force-time traces. To estimate the unfolding step size, the DNA extension estimated by the eWLC fit was subtracted from the total extension. The step size was estimated from the jump of the net extension over time. Limited by the noise, the minimum step size we could estimate is on the level of ∼5 nm. The unfolding force histogram was converted to the force-dependent folded state lifetime based on the previous work ^42^ which included the correction for the force-dependent loading rate resulted from the compliance of the DNA linkers. All optical tweezer data analysis was performed using MATLAB software.

### Motor pulling assay

The flow channel was prepared as aforementioned using 0.01 mg/mL anti-digoxigenin antibody (Roche) and passivated with 1% F127 and 2 mg/mL casein. To prepare the GMPCPP-capped microtubule, digoxigenin-labeled GMPCPP seeds were first polymerized (4 μM tubulin, 0.4 μM digoxigenin-tubulin, 1 mM MgCl_2_, 1 mM GMPCPP) at 37 °C for 30 min. The GMPCPP seeds were centrifuged and resuspended in BRB80 at a tubulin concentration of ∼4 μM (assuming ∼70% recovery). The DNA-tubulin mix containing 20 μM of oligo-tubulin (labeling density ∼3%), 2 μM of digoxigenin-tubulin, ∼8 nM 3.8 kb biotinylated DNA linker, 6 mM MgCl_2_, 1.5 mM GTP were incubated on ice for 15 min. The DNA-tubulin mix was then quickly warmed up to 37 °C and mixed with the GMPCPP-seeds in a 2:1 volume ratio. The microtubules were then polymerized at 37 °C for 25 min and quickly diluted with 10 times volume of BRB80-taxol solution (BRB80 containing 10 μM taxol). The taxol-stabilized microtubules were centrifuged and resuspended in BRB80-taxol to remove any unpolymerized tubulin and DNA molecules. The microtubules were then introduced into the flow channel; following attachment to the surface, the channel was washed with BRB80-taxol. To perform the capping, taxol was quickly washed out by flowing in BRB80 followed by the GMPCPP-tubulin capping mix (4 μM of unlabeled tubulin, 0.05 mg/mL neutravidin, 5 mM DTT, 0.5 mM GMPCPP, 0.1% Tween20, 2 mg/mL casein). The capping was performed at 28 °C for ∼5 minutes and the channel was washed with BRB80 again. This taxol-washout capping strategy allowed us to bind the GDP-microtubule segment to the surface and ensure that all DNA-tubulin subunits contained GDP rather than GMPCPP. Note that these microtubules should be free of taxol during the motor pulling experiments because the mean unbinding time of taxol is less than 10 s ^71^, and we also observed occasional shrinkage of GDP-microtubules during the course of our experiments.

To prepare biotinylated kinesin, rk430-mScarlet-SNAP (7 μM monomer) was incubated with benzylguanine-biotin (NEB) in a 4 to 1 molar ratio at room temperature for 15 minutes, which we found to be sufficient for complete reaction. This labeling ratio reduced the probability that a single kinesin dimer was labeled with two biotin moieties and only half of the kinesin dimers contained biotin (i.e. 50% labeling density). The biotinylated kinesin was stored on ice until the microscopy experiments. To visualize and stretch the DNA-linkers conjugated to GDP-microtubules, various concentrations of biotinylated kinesin were added into the oxygen scavenger solution containing ATP and SYTOX-Green (40 mM glucose, 0.04 mg/mL glucose oxidase, 0.02 mg/mL catalase, 0.2 mg/mL casein, 10 mM DTT, 2 mM Trolox, 0.1%Tween 20, 1 mM ATP, 40 nM SYTOX-Green) and flowed into the flow channel. The DNA molecules were then immediately visualized by TIRF microscopy with a 0.5 s time interval (or 1 min/frame for the photodamage controls). All experiments were performed at 28 °C.

All analyses of the motor pulling assay were performed by using Fiji ^72^. Rupture time was measured from kymographs. A sharp decrease of stretching velocity typically occurred when the DNA linkers were fully stretched. The rupture time was determined as the time from this velocity transition (with verification by checking whether the DNA was stretched close to its contour length) until the rupture events that corresponded to the removal of GDP-tubulin. Traces that showed clear photocleavage events were excluded from the rupture time measurement.

## Author contribution

Y.-W.K., M.M. and J.H. conceived the project; Y.-W.K. performed all experiments and data analysis with the assistance of M.M. and J.H.; preliminary data was collected by Y.-W. K. and M.M.; M.M., Y.-W.K. and Y.T. set up the optical tweezer instrument; Y.-W.K. and J.H. wrote the paper with the input of all authors.

## Acknowledgement

We thank Dr Veikko Geyer, Dr Maijia Liao and Dr Anna Luchniak for the helpful discussion. We thank Dr Ziad Ganim, Dr Ya-Na Chen and Dr Nandan Pandit for their help in the development of optical tweezer assay, and the Yale Keck Proteomics Center for the mass spectrometry analysis. We thank Dr Tom Pollard, Dr Enrique De La Cruz and Dr Yongli Zhang for the feedback on the work. Y.T. acknowledges the support of the Alexander von Humboldt Foundation through the Feodor Lynen Research Fellowship. This work was supported by NIH Grant R01 GM139337 (to J.H.).

## Data Availability Statement

The data and code that support the findings of this study are available from the corresponding author upon request.

