## Supplementary information for "The force required to remove tubulin from the microtubule lattice"

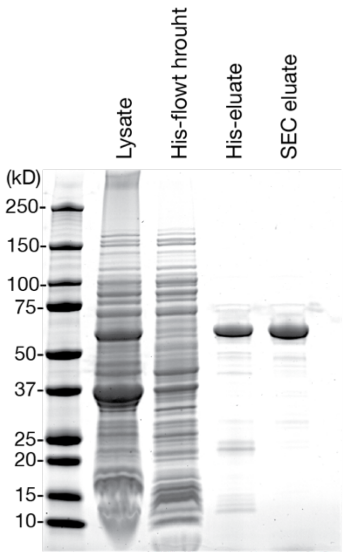


**Figure S1. SDS-PAGE of purification steps of recombinant tubulin tyrosine ligase (TTL).** Human TTL with N-terminal $His_6$-SUMO tag was expressed in E. coli and purified by HisTrap affinity column and size exclusion chromatography (SEC). Predicted molecular weight of $His_6$-SUMO-TTL: 57 kD.

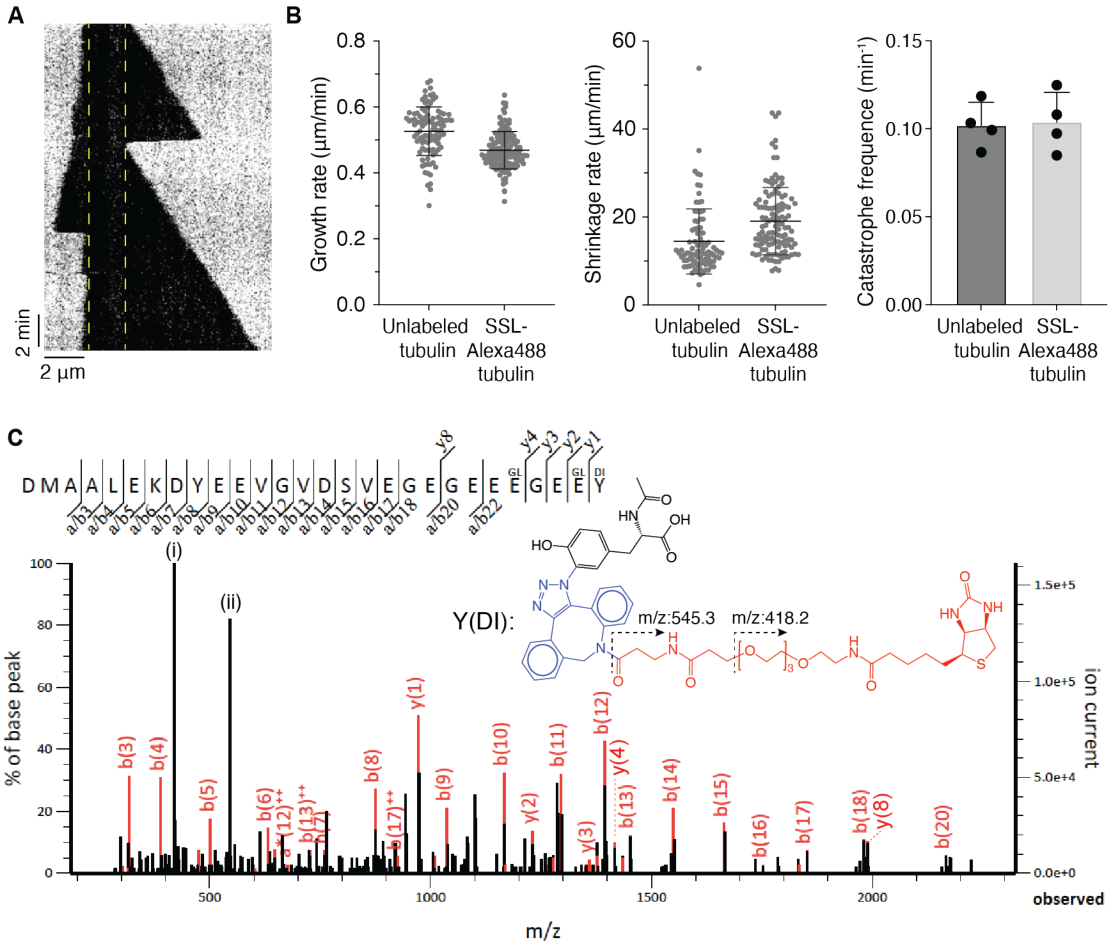


**Figure S2. TTL-dependent site-specific labeling had little effects on microtubule dynamics.** (A) Example kymograph of a dynamic microtubule extension growing off a GMP-CPP stabilized microtubule seed polymerized from 8 µM Alexa Fluor 488-conjugated tubulin using the TTL-dependent site-specific labeling (SSL) method imaged by IRM. The seed position is marked by yellow dashed lines. (B) Comparison of the dynamic properties of 8 µM unlabeled and SSL-Alexa Fluor 488-tubulin (labeling density 43%) imaged by IRM from four independent measurements. Dynamic parameters of unlabeled vs. SSL-tubulin (mean ± SD): growth rates (0.53 ± 0.07 vs. 0.47 ± 0.06 µm/min; *n* = 99 vs. 131 events); shrinkage rate (14 ± 7 vs. 19 ± 8 µm/min; *n* = 89 vs. 119 events); catastrophe frequency (0.102 ± 0.013 vs. 0.104 ± 0.008 $min^{-1}$; *n* = 4 experiments). (C) Example MS/MS spectrum of α-tubulin C-terminal peptide conjugated to biotin via the site-specific labeling method. Y(DI) shows the [3+2] cycloaddition product on the C-terminal tyrosine and was used as custom modification in the MASCOT search. Mono- and polyglutamate chains (GL) up to 4 glutamate residues were included as possible PTMs in the MASCOT search as well. The detected ion fragments corresponding to the cleavage of backbone bonds are highlighted in red. Note that two reporter ions (peak i and ii) corresponded to the fragmentation at the linker (dashed arrows).

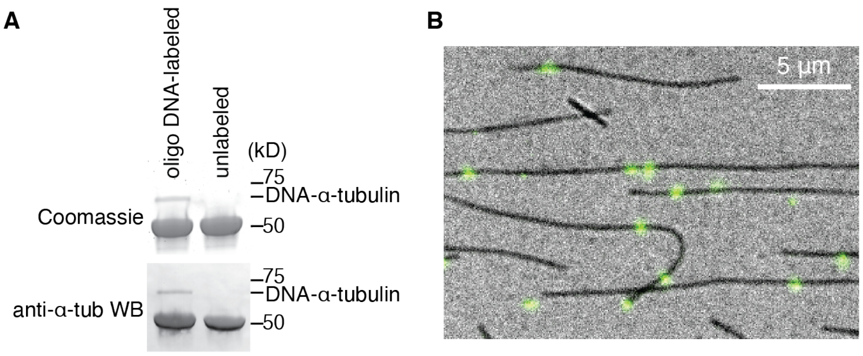


**Figure S3. Incorporation of oligo-DNA handle and long DNA linkers.** (A) Successful conjugation of oligo-DNA to α-tubulin via the site-specific labeling method was confirmed by the molecular weight shift of the oligo-DNA-labeled tubulin. (B) Example of GMPCPP-stabilized microtubule with long DNA-linkers (green puncta). The DNA linkers were stained with SYTOX Green and imaged by TIRF microscopy. The microtubules were imaged by IRM.

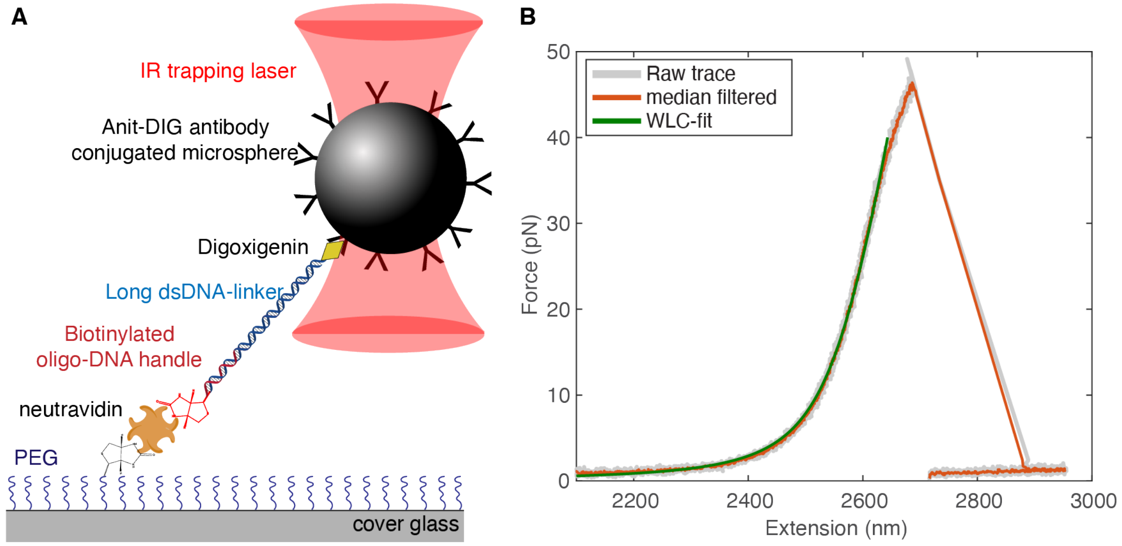


**Figure S4. Surface-tethered DNA linker pulling controls.** (A) Experimental scheme of the surface-tethered DNA control setup. The DNA linkers were first hybridized to the biotinylated oligo-DNA handle (identical sequence as the handle used in the tubulin pulling experiments). The coverslips were first covalently biotinylated followed by binding of neutravidin. The DNA-linkers hybridized to biotinylated DNA oligo were then anchored onto the surface via biotin-neutravidin binding. Note that the same DNA linker and microspheres were used in both the control experiment and the taxol-microtubule pulling experiments. (B) Example force extension curve of stretching surface-tethered DNA linker.

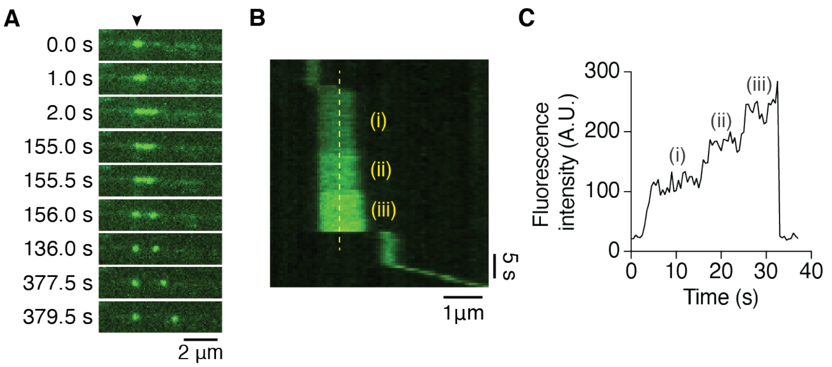


**Figure S5. Examples of photocleavage and step-wise force increase in the kinesin pulling assay.** (A) Example time-lapse images of a DNA photocleavage event after stretched due to photo-induced double-stranded break. The anchor point is indicated by the arrowhead. (B)(C) Example kymograph and fluorescence intensity line-scan (yellow dashed line in (B)) showing step-wise increase of DNA-fluorescence intensity during the stretching phase. Three distinct steps can be identified from the intensity trace in (C). The increase of intensity corresponded to the increase of pulling force on the DNA (King et al., 2018). The step-wise increase of force was likely due to the increase number of kinesin molecules engaging in the force generation process.

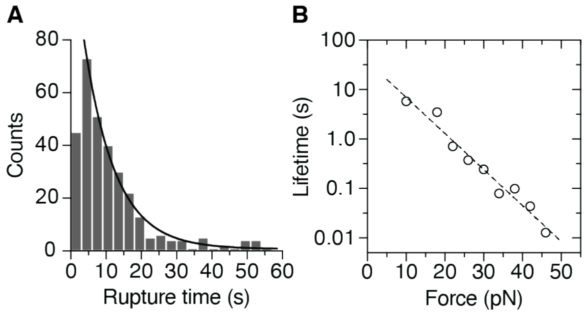


**Figure S6. Rupture time from kinesin pulling and optical tweezer assays. (A)** Rupture time histogram of kinesin-pulling assays (pooled from 4.5 to 9 nM; total 333 events; ~95% of data is within 0 to 60 s and was plotted here). The histogram was fitted with a single exponential decay (black line) with the first bin left out of the fit due to the difficulty of measuring short rupture time (< 3 s). Decay time of exponential fit was 8.8 s; $R^2$ = 0.988. **(B)** Force-dependent rupture time $\tau(F)$ was estimated from the rupture force histogram obtained in the taxol-stabilized microtubule pulling assay using optical tweezer in Fig. 3C by applying the method described in (Dudko et al., 2008). Fit with Bell model $\tau_0$ = 37.1 ± 1.5 s, $x^{\ddagger}$ = 0.69 ± 0.05 nm; $R^2$ = 0.96. Note that this is an underestimation of the true force-dependent rupture time of tubulin subunit since the DNA-bead rupture limit the maximum force that can be measured.

**Table S1. Primer pairs for DNA linker preparation**

| Primer 1 | Primer 2 | DNA linker size (kb) |
|---|---|---|
| 5'-TCTAAGTGACGGCTGCATACTAACC-3' (5' end and underlined bases were labeled with digoxigenin) | 5'-CGCCTGCGTAGGATATCGCAGATACCGCATCAGTCCAXCAACGGTCGATTGCCTGACGGA-3' (X: abasic) | 8.2 |
| 5'-biotin-GCCAATGCGCTTACTGATGCGG-3' | 5'-CGCCTGCGTAGGATATCGCAGATACCGCATCAGTCCAXGGTTTCACTGCTGGCGTATGACC(X: abasic) | 3.8 |

### Supplementary Information References:


1. Dudko, O.K., Hummer, G., and Szabo, A. (2008). Theory, analysis, and interpretation of single-molecule force spectroscopy experiments. Proc. Natl. Acad. Sci. *105*, 15755–15760.
2. King, G.A., Biebricher, A.S., Heller, I., Peterman, E.J.G., and Wuite, G.J.L. (2018). Quantifying Local Molecular Tension Using Intercalated DNA Fluorescence. Nano Lett. *18*, 2274–2281.
